# FunOMIC: Pipeline with built-in Fungal Taxonomic and Functional Databases for Human Mycobiome Profiling

**DOI:** 10.1101/2022.05.09.491154

**Authors:** Zixuan Xie, Chaysavanh Manichanh

## Abstract

While analysis of the bacterial microbiome has become routine, that of the fungal microbiome is still hampered by the lack of robust databases and bioinformatic pipelines. Here, we present FunOMIC, a pipeline with built-in taxonomic (1.6 million marker genes) and functional (3.4 million non-redundant fungal proteins) databases for the identification of fungi. Applied to more than 2,600 human metagenomic samples, the tool revealed fungal species associated with geography, body sites, and diseases. Correlation network analysis provided new insights into inter-kingdom interactions. With this pipeline and two of the most comprehensive fungal databases, we foresee a fast-growing resource for mycobiome studies.

## Introduction

Fungi ubiquitously exist as commensals in various body sites of humans, including the gastrointestinal tract (GIT), oral cavity, vagina, and skin (Seed 2014). Under certain circumstances, some of these fungal commensals, identified as pathobionts, could cause harm (Seed 2014; Liguori et al. 2016). Also, bacterial-fungal interactions have been reported to exacerbate, reduce, or resist disease caused by fungal infection (van Tilburg Bernardes et al. 2020; Santus et al. 2021). The colonised fungi are highly variable across populations (Sun et al. 2021), which may prevent the establishment and, thereby, the identification of key players among the fungal community in humans. It is therefore critical to investigate commensal fungi and their interactions with the host and commensal bacteria in a large-scale study.

Unlike the prokaryotic community in the human microbiome, the fungal population, known as mycobiome, is still understudied due to various reasons, including the challenge associated with unculturable microorganisms, the extremely low abundance among the human microbiome community (Qin et al. 2010), inter-individual variability, and the lack of a comprehensive database. Over the last decades, along with the rapid development of high-throughput sequencing (HTS) technology, the study of the human bacterial and fungal microbiome has gradually moved from culture-dependent towards culture-independent methods (Seed 2014).

The characterization of the mycobiome has been catalysed by targeted HTS of the internal transcribed spacer (ITS) or the 18S rRNA (18S) region located inside the ribosomal region. Similar to the 16S rRNA (16S) gene in prokaryotes, the ITS and 18S regions have conserved and highly variable segments among different fungal organisms. Moreover, the ITS has been recognised as a universal DNA barcode marker for fungi (Schoch et al. 2012). The current knowledge of human mycobiome derives mostly from the analysis of ITS and 18S amplicon sequencing (Andersen et al. 2013; Del Campo et al. 2019). However, as for the 16S amplicon sequencing approach (Louca et al. 2018), ITS and 18S approaches can introduce biases due to variability in amplification efficiency (Engelbrektson et al. 2010), problems related to species delineation, and the large variations in gene copy numbers, which limits the relative abundance analysis between closely related species (Lofgren et al. 2019). As an alternative to ribosomal DNAs, a set of single-copy marker genes can be candidates for taxonomically annotating the microbiome. They have been shown to provide higher resolution than 16S in prokaryotic species delineation (Mende et al. 2013) and have been used to estimate relative abundances and richness of bacterial members in human faecal microbiomes.

With the decreasing cost of sequencing, the shotgun approach, which can capture more unbiased information from the gene pool of microbial genomes within an environment than the amplicon approach, has emerged as a more attractive tool in microbiome research. Various strategies and databases have been developed to determine eukaryotic community compositions from metagenomic data (West et al. 2018; Marcelino et al. 2020; Lind and Pollard 2021), yet few of them tackled fungi in the context of the human microbiome.

To enable a more precise analysis of the human mycobiome, we propose herein two built-in fungal databases, FunOMIC-T and FunOMIC-P, integrated into an automated pipeline for taxonomic and functional profiling, respectively. The functionality of the pipeline is achieved by mapping next-generation sequencing reads to the two FunOMIC databases. FunOMIC-T contains more than 1.6 million single-copy marker genes from 4,839 high-quality fungal genome data. FunOMIC-P includes more than 3 million fungal proteins, being an integration of the corresponding coding genes of the collected fungal genomes with the fungal subset of the Uniprot database. FunOMIC was used to analyse a publicly accessible set of 2,679 human metagenome samples, which revealed fungal taxonomic and functional signatures associated with clinical and demographic metadata.

## Results

### Characteristics of the taxonomic and functional FunOMIC database

To build a database for taxonomic profiling of environmental fungal species, more than 1.6 million fungal single-copy marker genes were extracted from 4,816 fungal high-quality genomes and draft genomes by aligning them to a set of 758 fungal universal orthologs from OrthoDB (Fig. 1). The newly constructed database, FunOMIC-T, covers eight fungal phyla, among which three (Ascomycota, Basidiomycota, and Mucoromycota) represented more than 98% of the genomes (Fig. 1A). At lower taxonomic levels, they encompassed 475 genera, 1,916 species, and 4,537 strains.

**Fig 1.**
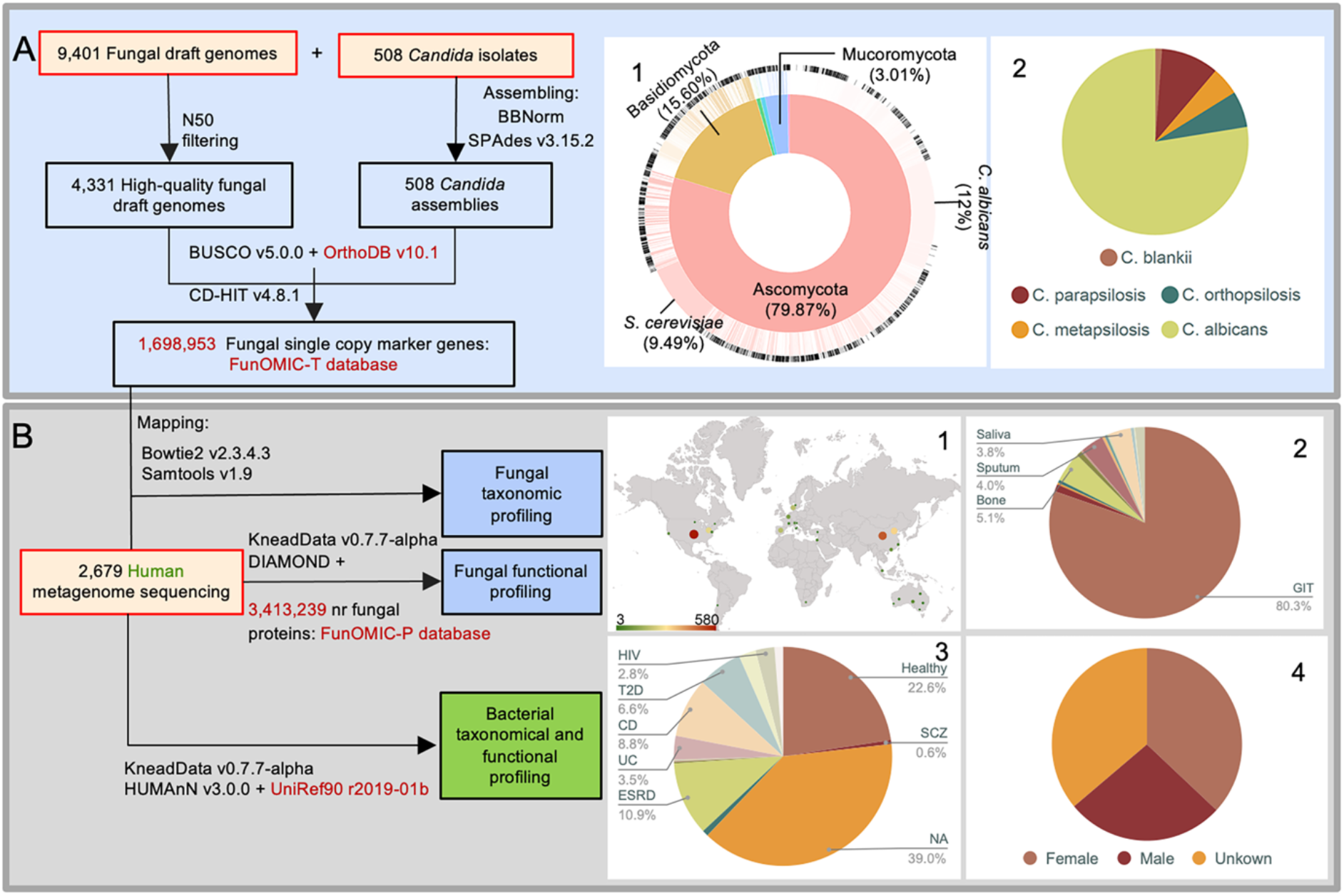
Workflow of the construction of the FunOMIC database and its application in metagenomic analysis. **(A)** Recovery of fungal single-copy marker genes from fungal draft genomes and *Candida* isolate sequencing reads downloaded from NCBI and JGI. A.1) Distribution of the fungal draft genomes at the phylum and species levels in FunOMIC-T (Taxonomy). A.2) Distribution of *Candida* assemblies at the species level. **(B)** Fungal and bacterial taxonomic and functional profiling of the 2,679 metagenomic datasets downloaded from NCBI. B.1) Geographical location of the collected human metagenomes. B.2) Proportions of the collected human metagenomes by body sites. B.3) Proportions of human metagenomes by disease type (HIV=human immunodeficiency virus; T2D=type 2 diabetes; CD=Crohn’s disease; UC=ulcerative colitis; ESRD=end-stage renal disease; SCZ=schizophrenia). B.4) Distribution of the collected human metagenomes by gender.

It has been reported that 99.9% of human metagenome sequences are from bacteria(Qin et al. 2010) and that, bacterial sequences are ubiquitous in eukaryotic genomes (Lind and Pollard 2021). Validation of the absence of bacterial sequence contamination in the fungal database is, therefore, critical. To address this requirement, the FunOMIC-T database was mapped to the UHGG dataset, which contains 204,938 non-redundant genomes from 4,644 gut prokaryotes (Almeida et al. 2021). Only less than 0.01% of the fungal marker genes mapped to the UHGG, demonstrating that this fungal taxonomic database was specific enough to detect mostly fungal sequences.

A bacterial environmental mock community was also created. For this, we collected 903 genomes from 458 bacterial species found to inhabit human bodies (Supplemental Table S1). These genomes were then simulated into 19,301,201 Illumina formatted sequencing output reads and mapped to the FunOMIC-T database. The mapping rate of this artificial community to the database was also less than 0.01%. Lastly, a mixed mock community was also created comprising the top 20 bacterial species and top 20 fungal species identified during the taxonomic profiling of the metagenomes (Supplemental Table S1). To better mimic real human metagenomes, the ratio of the number of simulated bacterial reads over fungal reads was set to nearly 1,000 (999021 bacterial reads and 1,046 fungal reads) (Qin et al. 2010). As expected, none of the 999,021 bacterial reads aligned against FunOMIC-T, leading to a specificity (false positive / (false positive + true negative)) of 0.9999.

Given the numerically small proportion of fungal sequences in human metagenomes, the fungal functional analysis was not relevant in almost all the published human mycobiome studies. To address this knowledge gap, in the present work, we also proposed a protein database specifically for environmental fungal functional profiling. The FunOMIC-P database consists of 3,413,239 non-redundant fungal protein sequences integrated from NCBI, JGI, and UniProt (see Methods section above, Fig. 1B). Evaluation and validation were also performed by a mixed mock community constituted by the top species mentioned above. The available coding gene sequences of these species were simulated into 439,798 Illumina formatted sequencing output reads and mapped to the FunOMIC-P database. We tuned the Diamond blastx function with nine different combinations of parameters to optimize mapping performance. With the threshold of read coverage > 95%, identity percentage > 99%, and an e-value < 10e-10, we obtained the highest mapping rate of the fungal reads, where around 70% of the hits passed this threshold. More than 50% of the mis-mapped bacterial genes were related to ATP synthase (Supplemental Table S1).

### Characteristics of the 2, 679 metagenomes

A set of 2,679 metagenomes, which encompassed a total of 9077.12 Gb, collected from 27 bioprojects are listed in Supplemental Table S2. Taxonomic profiling of the metagenomes against FunOMIC-T detected fungal DNA sequences in 1,950 metagenomes (72.9%) which was much higher than the ratio reported in previous shotgun sequencing studies analysing the human mycobiome. Lind *et al*., reported a detection rate of less than 20% and Olm *et al*., found 6% in their cohorts (infant). The 1,950 metagenomes were collected from 14 countries, 12 body sites, and 19 health and disease conditions (Table 1). The average mapping rate was 4.72E-05 (8.16E-09 min, 1.1E-02 max).

**Table 1.**
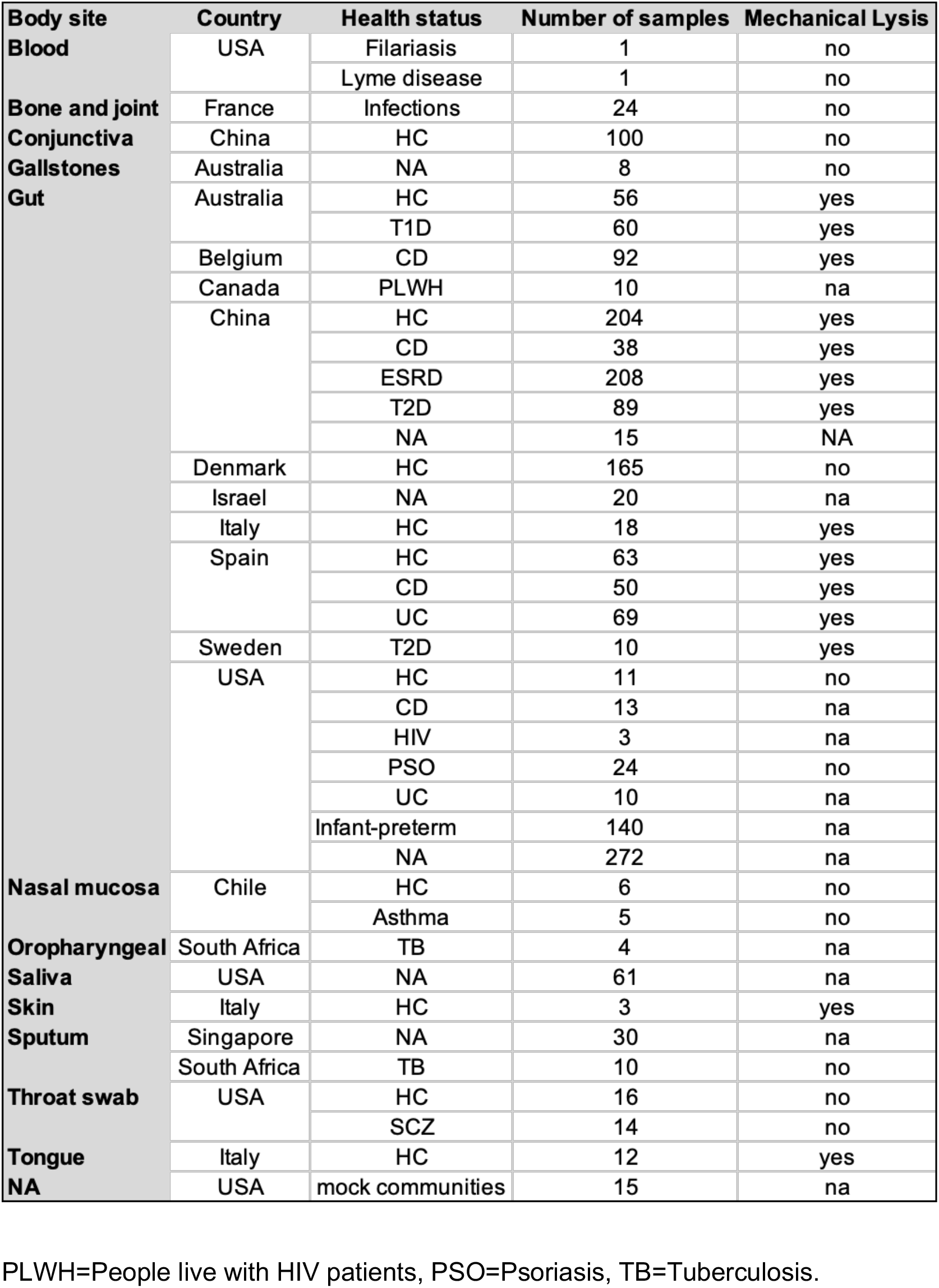
Summary of the characteristics of the 1,950 human metagenomes.

Gut samples comprised the majority of the dataset (84%), followed by conjunctiva (5%), saliva (3%), and throat swab (1.5%). Among the diseases evaluated, Crohn’s disease (CD), ulcerative colitis (UC), end-stage renal disease (ESRD), type 1 diabetes (T1D), and type 2 diabetes (T2D) accounted for 779 faecal samples, whereas 500 faecal samples were obtained from healthy individuals.

All biological specimens were extracted by at least 10 different protocols, for which mechanical lysis, previously reported as a crucial step during the DNA extraction process to recover an optimum microbial diversity (Costea et al. 2017), was applied in 1,049 samples (53.8%).

### Fungal community structure, diversity, and functions of the 1950 metagenomes

Five phyla, 232 genera and 475 species were identified in the 1,950 metagenomes. More than 80% of the sequences were represented by two phyla (Ascomycota and Basidiomycota), two genera (*Saccharomyces, Candida*), and three species (*Saccharomyces cerevisiae, Candida albicans, Malassezia restricta*) (Fig. 2). Under healthy conditions, the gut mycobiome was dominated, in terms of relative abundance, by *Saccharomyces cerevisiae*, which was detected in 52.4% of the samples, while *Dacryopinax primogenitus* was found in 23.6%, *Yarrowia lipolytica* in 13.6%, and *Candida parapsilosis* in 11% of the samples. *C. albicans*, known as an opportunistic pathogenic yeast (d’Enfert et al. 2021), was found in only 4% of the GI tract samples of healthy individuals. The fungal species profiling data can be found in Supplemental Table S3. *Malassezia* predominated conjunctiva samples, whereas *Aspergillus* predominated the saliva mycobiome.

**Fig 2.**
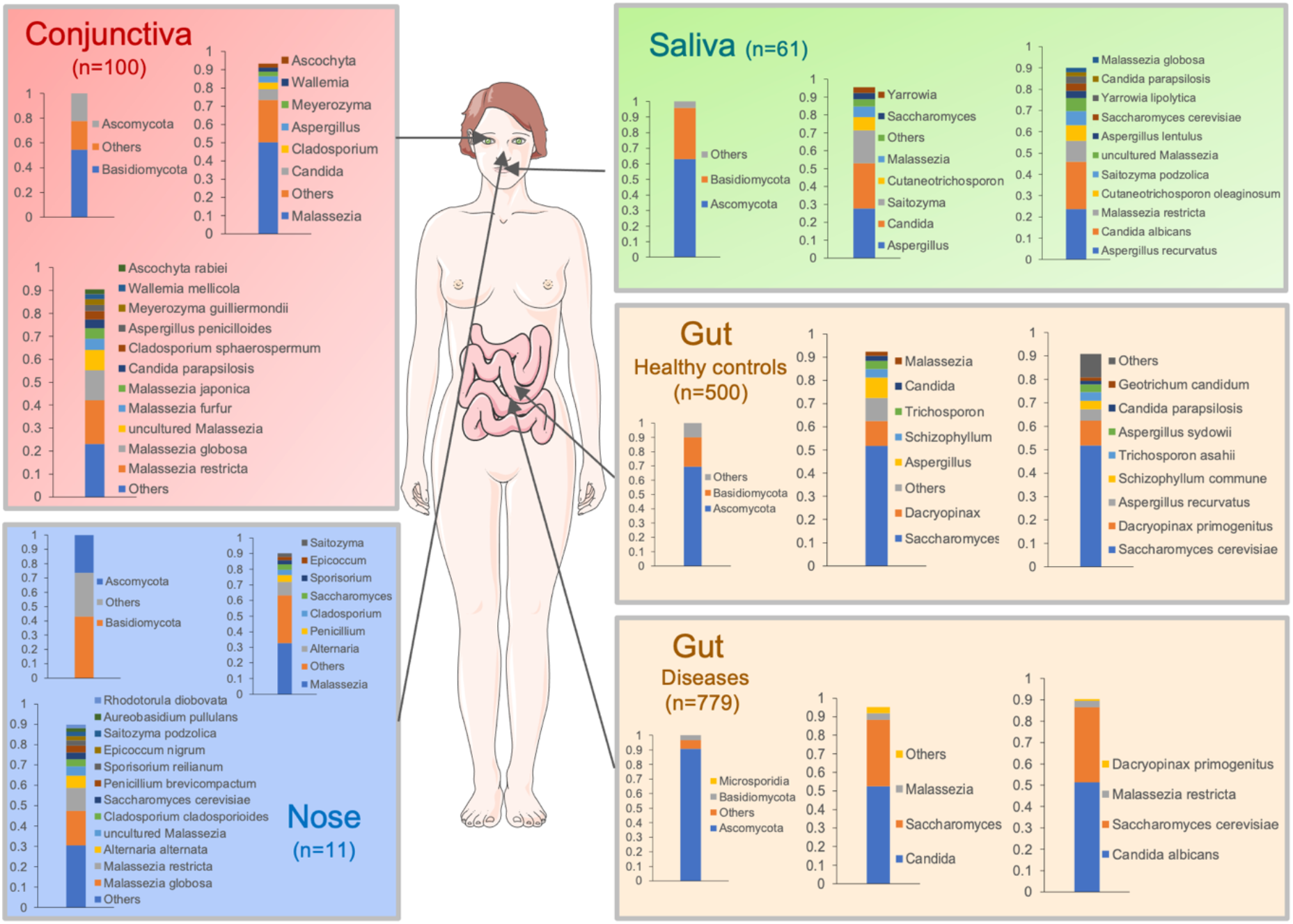
Fungal taxonomic profiling of several human body sites based on the 1950 shotgun metagenomic data using the FunOMIC-T database. Taxonomic profiling is displayed at the phylum, genus, and species levels. Only the mean relative abundance of the genera and species summing 90% of the sequence data is exhibited. Gut taxonomic profiling was performed for diseases including Crohn’s disease (CD, n=193; from the USA, Europe, and Asia), ulcerative colitis (UC, n=79 from Europe and the USA), end-stage renal disease (ESRD, n=208, from Asia), type 1 diabetes (T1D, n=60 from Australia), and type 2 diabetes (T2D, n=99 from Asia). 468 faecal samples did not have health status information in the metadata files. Health status and geo-localization of conjunctiva, nasal, and saliva samples are described in Table 1.

The number of observed fungal species in the 1,950 metagenomic samples ranged from 1 to 40 (median of 2), Chao1 index (Chao 1984) varied between 1 and 76.1 (median of 3), and the Shannon index (Shannon 1948) ranged from 0 to 3.36 (median of 0.62) (Supplemental Table S4). These three measurements indicated that the fungal community in humans is, in general, of very low diversity compared with the bacterial community, which could reach an average of 70 in terms of the Chao1 index (Serrano-Gomez et al. 2021).

While fungal taxonomic profiling of human microbial communities has increased considerably over the last 10 years through the sequencing of phylogenetic marker genes such as ITS2/18S, the fungal community function was scarcely investigated mainly due to, again, the lack of a comprehensive database. Using FunOMIC-P, we annotated the sequencing reads of the 1,950 human metagenomes using the DIAMOND aligner. In total 1,948 metagenomes successfully mapped to the database, the average mapping rate was 0.088% (5.42E-04% min, 1.2% max), consistent with that previously reported in Qin *et al*., for eukaryotic DNA (Qin et al. 2010).

Sixteen pathway classes and 120 pathways were detected from the metagenomes. Five pathway classes (Amino Acid Metabolism, Carbohydrate Metabolism, Nucleotide Metabolism, Energy Metabolism, Metabolism of Cofactors and Vitamins) and 29 pathways (Supplemental Table S5), along with unidentified pathways and pathway classes represented more than 80% of the sequences. The pattern of fungal functional structure indicated higher evenness compared with fungal taxonomic structure, i.e., the relative abundances of the pathways are closer instead of being dominated by one or two pathways.

### Association between metadata and mycobiome composition and functions

Next, we evaluated the contribution of available variables, collected from the metadata files, to the mycobiome composition variations using the adonis2 function from the vegan R package (Fig. 3). These variables included countries, health status, body sites, ages, gender, and bead-beating. Individually, countries and health status were the factors that contributed most to fungal composition and function variations; body sites and the bead-beating step also contributed to these variations, but to a lesser extent (FDR< 0.01, Fig. 3).

**Fig 3.**
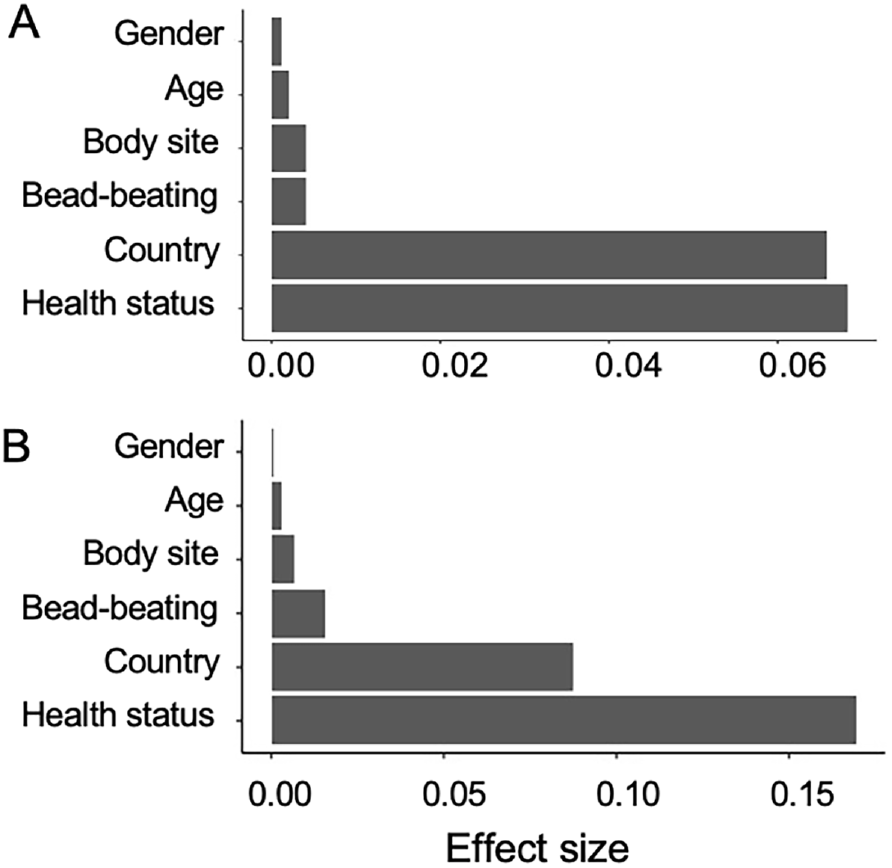
Effect size of variables on the mycobiome community. The impact of the covariates on mycobiome composition (A) and function (B) was tested by performing a univariate analysis (adonis2) on the 1,950 metagenomes. The effect was considered significant when FDR < 0.05.

Associations between these variables and individual taxa were then examined using generalised linear models implemented in the MaAsLin2 (Microbiome Multivariable with Linear Models) package. Five fungal species (*Aspergillus recurvatus, Malassezia restricta, Saccharomyces cerevisiae, uncultured Malassezia, Yarrowia lipolytica*), which were among the 10 most prevalent and abundant fungal species (Supplemental Table S3), were found associated with health status, country, and body sites (Supplemental Fig. S1A). This finding suggests that the high variability of the human mycobiome could be linked to these five species. Interestingly, *Yarrowia lipolytica* was found positively associated with bead-beating (Supplemental Fig. S1B), which could be explained by its relatively higher fraction of chitin (10.3-18.9%) in the cell wall compared with *S. cerevisiae, C. albicans*, and *M. restricta* (Chattaway et al. 1968; Chaffin et al. 1998; Stalhberger et al. 2014).

We found that geography, health status, and body sites had marked effects on the variability of most of the fungal pathway classes among the 16 that we recovered from all samples, yet bead-beating did not impact the compositions of fungal pathways, as reported for fungal taxa (Supplemental Fig. S1).

**Supplemental Fig. S1.**
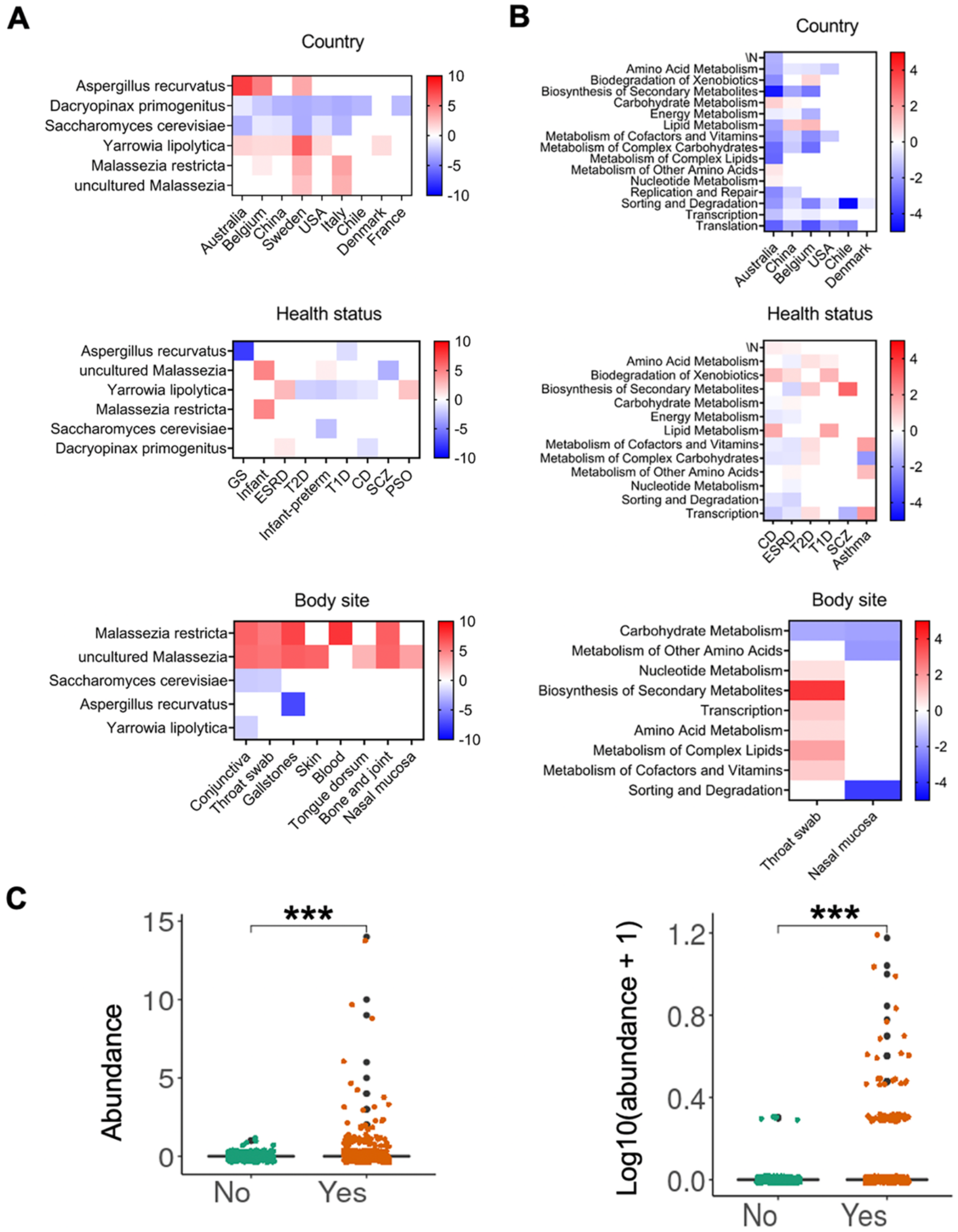
Fungal species and pathway classes in different groups of mycobiomes. Fungal species **(A)** and pathway classes **(B)** associated with different countries, health status and body sites. The colours represent the sign of the association (red means positive, blue means negative). The intensity of the colours represents the degree of association (darker means stronger association). Spain, healthy and gut were set as the reference level for each of the models, respectively. **(C)** Significant differences in *Yarrowia lipolytica* abundances or log10 transformed abundances between bead-beaten samples and non-bead beaten samples.

### Core taxonomic fungal microbiomes of different body sites and different countries

To identify groups of key taxa that may influence the microbiome community, we applied the concept of core microbiome across body sites and geography, taking into account health status. For this purpose, fungal species with an occurrence of over 50% in the respective set of metagenomes of interest, in which fungi were detected, were defined as the core mycobiome. The 50% occurrence threshold was chosen based on the review of the core bacterial microbiome published by Neu *et al*. (Neu et al. 2021), but an abundance cut-off was not applied to avoid missing any lowly abundant fungal species. We summarised the core mycobiome for body sites (Table 2) and countries (Table 3). In the human gut mycobiome of non-infants, *S. cerevisiae* was found to be the only member of the core gut mycobiome, except for CD and T1D patients who were dominated by *Aspergillus recurvatus*. The core gut mycobiome of infants consisted of only species from the *Malassezia* genera, in accordance with several previous studies (Boutin et al. 2021; Ventin-Holmberg et al. 2021). In other body sites, except saliva, several *Malassezia* species were the most detected members of the core mycobiome. The saliva mycobiome was driven by *Aspergillus recurvatus*.

**Table 2.** Core fungal species of different body sites.

| Bodysite | Health status | Core fungal species (>50% prevalence) |
| --- | --- | --- |
| Gut | HC (n=262) | *Saccharomyces cerevisiae* |
|  | CD (n=109) | *Aspergillus recurvatus* |
|  | ESRD (n=106) | *Saccharomyces cerevisiae* |
|  | UC (n=55) | *Saccharomyces cerevisiae* |
|  | T1D (n=40) | *Aspergillus recurvatus* |
|  | T2D (n=50) | *Saccharomyces cerevisiae* |
|  | PSO (n=16) | *Saccharomyces cerevisiae* |
|  | PLWH (n=7) | *Saccharomyces cerevisiae* |
|  | Infant (n=14) | *Malassezia globosa*, *Malassezia restricta*, uncultured *Malassezia* |
| Nasal mucosa | HC (n=5) | *Alternaria alternata*, *Malassezia globosa* |
| Conjunctiva | HC (n=76) | *Malassezia globosa*, *Malassezia restricta*, uncultured *Malassezia* |
| Saliva | NA (n=38) | *Aspergillus recurvatus* |
| Throat swab | HC (n=14) | *Schizophyllum commune*, *Malassezia restricta*, uncultured *Malassezia* |
|  | SCZ (n=12) | *Candida albicans*, *Malassezia restricta* |
| Tongue dorsum | Infant (n=8) | *Malassezia globosa*, *Malassezia restricta*, uncultured *Malassezia* |
| Bones and joints | BJIs (n=24) | *Malassezia globosa*, *Malassezia restricta*, uncultured *Malassezia* |
| Gallstone | GS (n=8) | *Malassezia globosa*, *Malassezia restricta*, uncultured *Malassezia* |

**Table 3.** Core fungal species of different countries.

| Country | Health status | Core fungal species (>50% prevalence) |
| --- | --- | --- |
| Australia | HC (n=46) | *Aspergillus recurvatus* |
|  | T1D (n=40) | *Aspergillus recurvatus* |
| Belgium | CD (n=76) | *Aspergillus recurvatus*, *Saccharomyces cerevisiae* |
| China | ESRD (n=106) | *Yarrowia lipolytica* |
|  | T2D (n=49) | *Saccharomyces cerevisiae* |
| Canada | PLWH (n=7) | *Saccharomyces cerevisiae* |
| Denmark | HC (n=118) | *Saccharomyces cerevisiae* |
| Italy | Infant (n=14) | *Malassezia globosa*, *Malassezia restricta*, uncultured *Malassezia* |
| Spain | HC (n=57) | *Dacryopinax primogenitus*, *Saccharomyces cerevisiae* |
|  | CD (n=38) | *Saccharomyces cerevisiae* |
|  | UC (n=52) | *Saccharomyces cerevisiae* |
| USA | PSO (n=16) | *Saccharomyces cerevisiae* |

Given that geographical difference contributes the most to fungal taxonomic structure variations, we also defined the core mycobiome for gut samples collected in different countries. We focused only on gut samples, as they represented the most available samples. *S. cerevisiae* appeared as a member of the core gut mycobiome in most countries (Table 3), which is in agreement with the aforementioned core mycobiome (Table 2). *A. recurvatus* was the only core fungal species among all the gut samples with different health status collected from Australia, whereas *Y. lipolytica* was that of the gut samples collected from end-stage renal disease (ESRD) patients in China (Table 3).

Core biochemical pathways, defined as pathways that have occurrences over 99% among all the samples with a relative abundance of over 1% (Neu et al. 2021), were also summarised for each body site and country with different health status (Supplemental Table S6). For countries, only gut samples, as the most available sample type, were considered. The majority of core fungal pathways were related to nucleotides, amino acids, energy and carbohydrate metabolisms, which are essential functions, indicating that the functionality of the human mycobiome is maintained across body niches and populations.

### Bacterial and fungal microbiome interaction

Next, we sought to evaluate the correlations between fungal and bacterial taxonomic composition in gut samples under healthy conditions, especially concentrating on core fungal species. Because of the failure in detecting the core mycobiome under healthy conditions from China, we focused on the healthy conditions of Denmark and Spain. To address this aim, we first performed a bacterial taxonomic and functional profiling of the metagenomic data. Due to a very extensive computational time requirement (6 hours/40 CPUs/sample on average), only a subset of 1,485 of the 2,679 metagenomic samples was processed (Fig. 1). We then carried out a correlation analysis with the SparCC correlation method, which handles compositional data (Friedman and Alm 2012) (34). In total, 4,184 significant (p < 0.05) inter-kingdom correlations were found in the Danish cohort, while 3,471 significant inter-kingdom correlations were found in the Spanish cohort, (Supplemental Table S7). In the Spanish cohort, the two core fungal species, *S. cerevisiae* and *D. primogenitus*, were found to correlate with the bacterial species *Haemophilus pittmaniae* positively and negatively, respectively (Fig. 4A). Beyond that, in the Spanish cohort, *C. albicans* was found to negatively correlate with *Megasphaera sp MJR8396C*, which was positively correlated with *D. primogenitus. C. albicans* was also found negatively correlated with *Lactobacillus sanfranciscensis, Bifidobacterium scardovii, Desulfovibrio fairfieldensis, Ruminococcus sp CAG563, Coprococcus catus*, and *Roseburia sp CAG309* (Supplemental Table S7, Fig. 4A), many of which are potential short-chain fatty acid (SCFA) producers (Parada Venegas et al. 2019). In the Danish cohort, significant correlations were found between the only core fungal species, *S. cerevisiae*, and seven bacterial species, of which five were negative (*Tropheryma whipplei, Prevotella sp CAG1124, Firmicutes bacterium CAG24, Gemella sanguinis*, and *Sutterella parvirubra*) and two were positive (*Bacteroides nordii* and *Prevotella stercorea*) (Fig. 4B).

**Fig 4.**
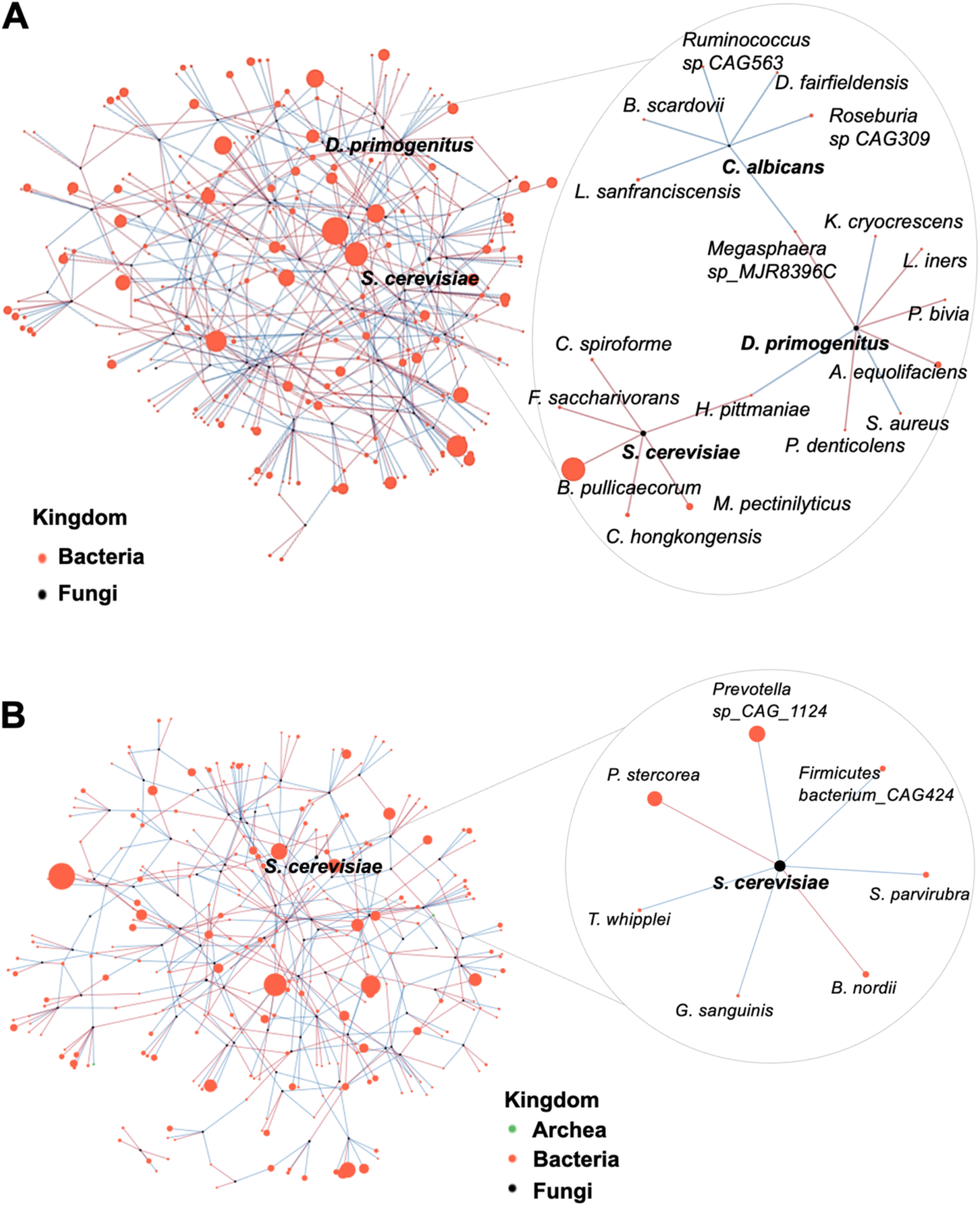
Interaction of fungal and bacterial communities in gut microbiome under healthy conditions. Correlation network between the relative abundance of fungal and bacterial species in the gut mycobiome under healthy conditions from Spain **(A)** and Denmark **(B)** using the SparCC algorithm. Each node represents a fungal/bacterial/archaeal species and their sizes are determined by relative abundances. The colours of the edges connecting two nodes represent positive (red) and negative (blue) correlations. For a better visual effect, only correlations with p-values less than 0.001 and an absolute correlation coefficient over 0.05 are represented.

We also applied SparCC to analysing correlations between fungal and bacterial functions in gut samples under healthy conditions. In the Danish cohort, 93 significant correlations were detected (Supplemental Table S7, Supplemental Fig. S2A), of which the strongest was the positive correlation (ρ=0.06, p<0.001) between the biosynthesis of secondary metabolites in fungi and the endocrine system in bacteria. In the Spanish cohort, 76 significant correlations were detected (Supplemental Fig. S2B), the strongest was a negative correlation (ρ=-0.13, p<0.001) between carbohydrate metabolism in fungi and signal transduction in bacteria. These functional inter-kingdom correlations could explain how bacteria and fungi interact in the microbiome community.

**Supplemental Fig. S2.**
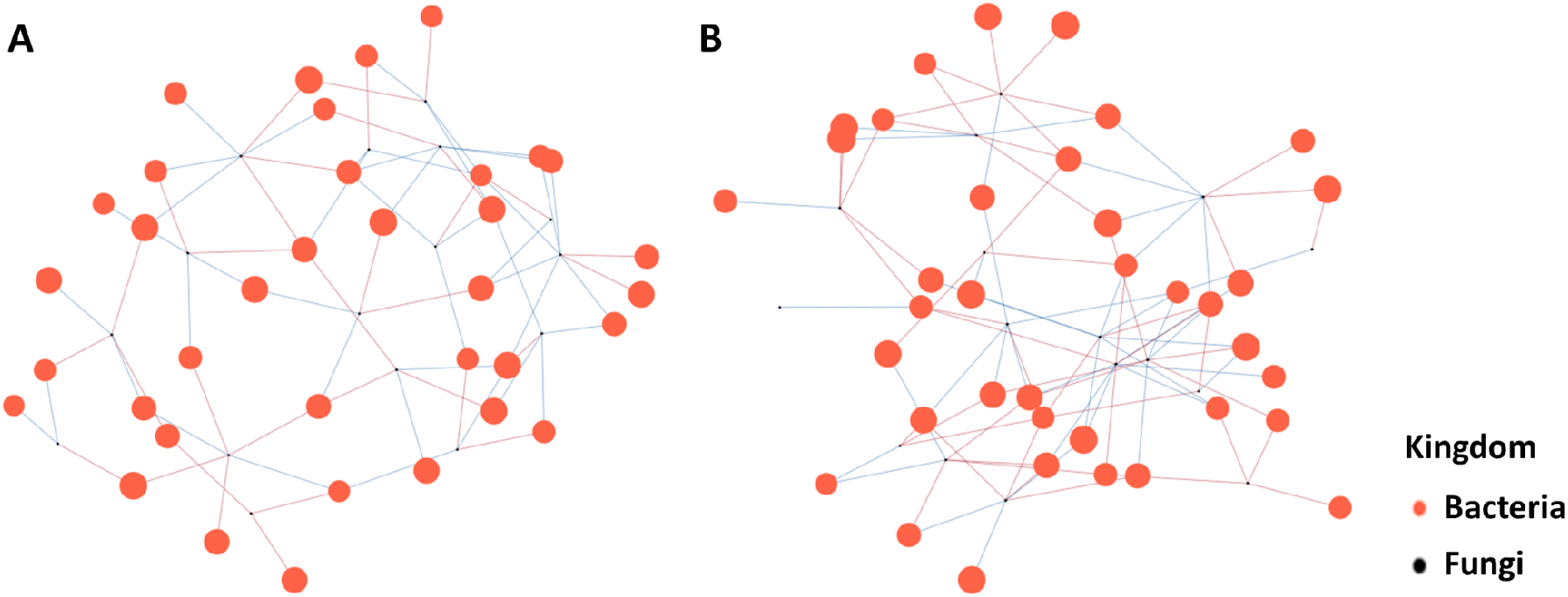
Interaction of fungal and bacterial functions in the gut microbiome of healthy individuals. Significant correlation (p<0.05) network between the relative abundance of fungal and bacterial functions in the gut mycobiome of healthy individuals from Spain **(A)** and Denmark **(B)** using SparCC algorithm. Each node represents a fungal/bacterial pathway class and their sizes are determined by the relative abundances. The colors of the edges connecting two nodes represent the positive (red) and negative (blue) correlations. For a better visual effect, only the correlations with p-values smaller than 0.001 and absolute correlation coefficient over 0.05 are represented.

## Discussion

Here, we have designed and validated FunOMIC, a metagenomic pipeline that integrates quality control, taxonomic profiling (FunOMIC-T), and functional profiling (FunOMIC-P) for a comprehensive analysis of fungi in environmental samples, and, particularly, in humans. First, to the best of our knowledge, FunOMIC offers the most comprehensive coverage of the reference fungal species and functions compared with other existing databases for profiling the human mycobiome. Indeed, FunOMIC-T, which contains more than 1.6 million fungal single-copy marker genes and covers 1,916 fungal species, exceeds the fungal spectrum of other similar tools (Donovan et al. 2018; Soverini et al. 2019; Lind and Pollard 2021). We also proposed FunOMIC-P which includes more than 3 million non-redundant fungal proteins, which is, to our knowledge, the first protein database proposed for analysing human mycobiome functions. Second, FunOMIC-T provided a smaller-sized taxonomic database with more accurate mapping possibilities for mycobiome profiling using universal conserved fungal genes instead of the full genome-based fungal reference database. Third, validations with different mock communities mimicking the human gut microbiome ensured extremely low bacterial read mis-mapping.

In this study, we applied the FunOMIC pipeline to a set of nearly 2,700 metagenomic human samples representing human microbiomes of different body sites from individuals with different health status and from different geographical regions. We corroborated previous human mycobiome results showing that the species *S. cerevisiae, C. albicans*, and *M. restricta* dominate the fungal communities in different human body sites (Ghannoum et al. 2010; Zhang et al. 2011; Hamad et al. 2017; Gupta et al. 2020). We found that geography and health status were the two most important factors contributing to the variabilities of human mycobiome taxonomic and functional compositions. Five fungal species (*A. recurvatus, M. restricta, S. cerevisiae, uncultured Malassezia, Y. lipolytica*) varied along with different countries, health status, and body sites. *C. albicans*, one of the most common human fungal pathogens (Kim and Sudbery 2011), negatively correlated with bacterial species that are mainly SCFA producers (Parada Venegas et al. 2019). This finding suggests that therapeutic strategies based on SCFA administration or on inducing SCFA producers could be implemented to control *C. Albicans* infection.

One important limitation of this pipeline is that the extraction and quality of single-copy marker genes rely on the completeness of the available fungal genomes, which may result in a lower coverage of fungal taxonomies compared with the fungal amplicon databases (Quast et al. 2013; Nilsson et al. 2019). Another limitation comes from the high inter-kingdom conservation of a portion of protein-coding genes. As a consequence, bacterial contamination was not totally preventable, even after applying an exceedingly strict mapping threshold to the fungal functional annotation. To overcome this drawback, filtration to remove the majority of bacterial reads before functional annotation could be included in the future update of this tool. Beyond that, in this study, FunOMIC was only applied to human microbiome data; in the future, applications with soil microbiome, marine microbiome, or other different environmental samples will be launched with FunOMIC to test its ability to handle other microbiome data.

## Conclusions

Taken together, our work presented here demonstrates that the proposed taxonomic database FunOMIC-T can effectively detect fungal species from shotgun metagenomic sequencing data. Together with FunOMIC-P, which to our knowledge, the first proposed functional database for mycobiome analysis, we believe that more mycobiome findings will be revealed in the future.

## Methods

### the aim, design and setting of the study

FunOMIC is a pipeline implemented with two fungal databases FunOMIC-T and FunOMIC-P aiming at providing automatic mycobiome analysis. Shotgun sequencing reads are directly mapped to the databases to obtain mycobiome taxonomic and functional profilings via the main program FunOMIC.sh. The main program and two databases can be downloaded from Manichanh Lab (vhir.org). Detailed establishment steps can be found below.

### Collection of fungal genomes

In total, 9,401 publicly available strain-level fungal genomes or draft genomes were downloaded from NCBI (https://www.ncbi.nlm.nih.gov/) and JGI MycoCosm (https://mycocosm.jgi.doe.gov/mycocosm/home) (Grigoriev et al. 2014) before January 25th, 2021. All fungal genomes with more than 500 contigs and N50 < 10 kbp were filtered out (Li et al. 2014), which led to a final set of 4,331 high-quality genomes and draft genomes. Genomic shotgun data from 508 *Candida* isolates were downloaded from 30 unique bioprojects from the NCBI SRA before February 4th, 2021 (https://www.ncbi.nlm.nih.gov/sra/). The accession numbers of the 4,839 combined reference fungal genomes are listed in Supplemental Table S8.

### Construction of the taxonomic and functional FunOMIC database

#### Identification of marker genes for establishing a taxonomic fungal database

Assembling *Candida* genomic sequencing reads was performed as described in the study of Montoliu-Nerin *et al*. (Montoliu-Nerin et al. 2020). Basically, each of the *Candida* genomic sequencing reads was normalised by BBNorm v38.9021 of BBtools (https://jgi.doe.gov/data-and-tools/bbtools/) with a target average depth of 100x. Then, normalised data were assembled by SPAdes v3.15.2 (Bankevich et al. 2012) (https://cab.spbu.ru/software/spades/). BUSCO (Benchmarking Universal Single-Copy Orthologs) version 5.0.0 (22) was used to identify marker genes using Fungi OrthoDB version 10.1 (Kriventseva et al. 2019) in the pool of 4,839 fungal genomes. BUSCO makes use of 758 HMMs (hidden Markov models) of fungal single-copy marker genes and was run using default parameters with the AUGUSTUS gene predictor (Manni et al. 2021). Genomes with less than 30 single-copy marker genes identified were discarded, resulting in a final set of 4,816 genomes. Clustering with a 99% identity threshold (Quast et al. 2013; Lind and Pollard 2021) was applied using CD-HIT (Fu et al. 2012) to remove redundancies, which led to a final set of 1.69 million fungal marker genes, referred here as FunOMIC-T.

#### Establishment of a functional fungal database

A protein database for fungal functional analysis was also constructed by collecting the corresponding amino-acid sequences that were available for 2,967 of the 4,331 genomes cited above and the 35,360 reviewed fungal proteins from UniProt (https://www.uniprot.org/), both before January 2022. Then, the proteins without an explicit annotation were discarded (1.5 million) leading to a total of 4.9 million genes. Redundancy was removed with a 95% identity clustering using CD-HIT (Qin et al. 2010). Finally, 3,413,239 non-redundant fungal proteins, referred to as FunOMIC-P, were obtained for fungal functional profiling. These protein accessions (from JGI, NCBI, UniProt) were then linked to EC numbers and KEGG pathways.

### Validation of the FunOMIC databases and the pipeline

To verify the absence of bacterial contamination (Lind and Pollard 2021) in the fungal database and to ensure specificity for fungal detection, we applied three different validation methods. Firstly, we mapped the 1.69 million fungal single-copy marker genes to the Unified Human Gastrointestinal Genome (UHGG), which is a gene catalogue that comprises 204,938 non-redundant genomes from 4,644 gut prokaryotes (Almeida et al. 2021) using bowtie2.

Because of the memory limitation of our computers (44 CPUs), we simulated sequencing reads of all the marker gene sequences (22 million paired-reads, 1-fold coverage, 11.2 GB out of 4.6 GB) to perform the alignment to the UHGG. Secondly, we simulated Illumina formatted sequencing output reads from a set of 903 bacterial genomes from 458 species that inhabit the human body collected from the NCBI to create a mock community for a bacterial community (Supplemental Table S1). The simulation was carried out by ART, a set of simulation tools that generate synthetic next-generation sequencing reads (Huang et al. 2012). The simulated reads were then aligned to FunOMIC-T. Thirdly, another mock community was created with the top 20 fungal species and top 20 bacterial species identified in the 2,679 human metagenomes collected (cited below). The genomes of these 40 species were used to simulate Illumina formatted sequencing output reads, which were then mapped to the constructed database. The lists of genomes used for creating the mock communities and the number of simulated reads can be found in Supplemental Table S1.

To validate the FunOMIC-P database, a mixed mock community was created with the available coding gene sequences of the aforementioned top fungal and bacterial species. Again, the coding gene sequences collected from NCBI were used to simulate Illumina formatted sequencing output reads, which were then mapped to the FunOMIC-P database using Diamond blastx function v2.0.8 with an e-value < 10e-10 to recover the fungal functional profiling. To optimise the alignment parameters, we tested nine different combinations using three different percentages of coverage (>90%, >95%, >99%) and three different percentages of identity (>90%, >95%, >99%).

### Collection of metagenomic data

We downloaded 2,679 public human shotgun metagenomic sequencing data from NCBI SRA before February 4th, 2021 (Leinonen et al. 2011) (https://www.ncbi.nlm.nih.gov/sra/). The 2,679-public human metagenomic data derive from 27 unique bioprojects, two of which were published in our previous studies (PRJNA514452, PRJEB1220). The metadata of all the human metagenomic data can be found in Supplemental Table S2. This metadata contains available information such as continent, country, city, latitude, longitude, sample source, gender, age, extraction procedure, and use of mechanical lysis during extraction.

### Aligning human metagenomic sequencing reads onto the FunOMIC database

After quality control and decontamination using KneadData v0.7.7-alpha (https://huttenhower.sph.harvard.edu/kneaddata/), Bowtie2 v2.3.4.3 was used to map the 2,679 metagenomic data to the FunOMIC-T database for fungal taxonomic annotation. Mapped reads were kept if more than 80% of the length aligned to the reference sequence with a q-score of over 30 (Qin et al. 2010; Milanese et al. 2019; Lind and Pollard 2021) by using Samtools v1.9. Diamond blastx function v2.0.8 was used to map the metagenomic data to the FunOMIC-P database (read coverage > 95% and identity percentage > 99% and e-value < 10e-10) for fungal functional annotation. An in-house script, which is freely available at our GitHub (https://github.com/ManichanhLab/FunOMIC), was used to recover the final fungal taxonomic and functional profiling.

### Prokaryotic taxonomic and functional profiling of human metagenomic data

After quality control and decontamination using KneadData v0.7.7-alpha (https://huttenhower.sph.harvard.edu/kneaddata/), we used MetaPhlAn v3.0.9 for profiling the composition of prokaryotic communities in the 2,679 human metagenomic data. Then, the HUMAnN v3.0 (Franzosa et al. 2018) (https://huttenhower.sph.harvard.edu/humann/) and the UniRef90 database (Suzek et al. 2015) were used to profile the abundance of prokaryotic metabolic pathways and other molecular functions.

### Statistical Analysis

All statistical analyses, except for SparCC correlation, were performed using R software 4.1.2 (2021-11-01). Alpha-and beta-diversity were calculated using the Phyloseq package. Beta-diversity was compared between different disease groups using the UniFrac distance metric with permutational multivariate analysis of variance (PERMANOVA) to identify significance (p ≤ 0.05). The associations between fungal profilings with variables from the metadata were measured using the MaAsLin2 package with age as the random effect (results were considered significant if FDR (false discovery rate) < 0.05). The correlations of taxonomic profilings or functional profilings between bacteria and fungi were performed using the Python script SparCC (Friedman and Alm 2012).

### Data access

The two built-in databases, FunOMIC-T and FunOMIC-P, are freely available at http://manichanh.vhir.org/funomic/. The source code of pipeline FunOMIC is freely available at our GitHub (https://github.com/ManichanhLab/FunOMIC).

## Abbreviations

CD: Crohn’s disease
ESRD: End-stage renal disease
FDR: False discovery rate
GS: Gallstones
HC: Healthy control
HTS: High throughput sequencing
ITS: internal transcribed spacer
NA: Not applicable
PLWH: People live with HIV
PSO: Psoriasis
SCFA: Short chain fatty acid
SCZ: Schizophrenia
TB: Tuberculosis
T1D: Type 1 diabetes
T2D: Type 2 diabetes
UC: Ulcerative colitis

## Authors’ contributions

CM obtained the funding; ZX built the databases and analyzed the data; CM and ZX interpreted the results and wrote the manuscript.

## Acknowledgements

Not applicable

